# Alternating 2D and 3D Culture Reduces Cell Size and Extends the Lifespan of Placenta-Derived MSCs

**DOI:** 10.1101/2025.05.20.655182

**Authors:** Ying Pan, Li Han, Yakun Yang, Xinran Wu, Aijun Wang, Liangqi Xie, Wuqiang Zhu, Shue Wang, Yuguo Lei

**Author notes:** authors contributed equally (Ying Pan, Li Han). Corresponding Authors: Yuguo Lei, Pennsylvania State University, PA, USA; Shue Wang, University of New Haven, CT, USA.

## Abstract

Mesenchymal stem cells (MSCs) offer significant therapeutic potential, but traditional 2D culture on rigid substrates results in progressive cell enlargement and senescence, reducing proliferative capacity and therapeutic potency. This poses a major challenge for widespread clinical application. We explored a novel strategy for placenta-derived MSCs combining 2D expansion with 3D spheroid culture to address these limitations. Our research shows that culturing MSCs as 3D spheroids significantly reduces individual cell size and size distribution compared to 2D culture. Spheroid formation is feasible in chemically defined media with minimal cell death, and is enhanced by extracellular matrix protein supplementation. Although MSCs do not proliferate in 3D suspension, an alternating 2D/3D culture protocol, transitioning cells between 2D flasks and 3D spheroids after each passage, effectively slows MSC enlargement and senescence over long periods. This alternating method also preserves MSC immunomodulatory function, unlike continuous 2D culture which leads to its loss. For scalability, we developed an RGD-functionalized alginate hydrogel tube (AlgTube) system that mimics this alternating environment, supporting both adherent growth and chemically triggered spheroid formation. This alternating 2D/3D culture strategy and the scalable AlgTube platform provide a foundation for developing next-generation MSC manufacturing technologies to meet future clinical demands.

## Introduction

Mesenchymal stem cells (MSCs) are multipotent stromal cells characterized by their capacity for self-renewal and differentiation into various mesenchymal lineages, including osteoblasts, chondrocytes, and adipocytes^1^. Originally identified in the bone marrow, MSCs have since been isolated from diverse tissues such as adipose tissue, umbilical cord, and dental pulp and placenta^1^. MSCs have garnered significant attention as therapeutics due to their exceptional safety profile and wide-ranging functions such as repairing tissues, promoting angiogenesis, inhibiting fibrosis, cytoprotection, anti-inflammation, neutralizing reactive oxygen species (ROS), inhibiting NETosis, suppressing T cell activation, promoting Treg differentiation, and polarizing M2 macrophages^1^.

For instance, MSCs have shown promise in reducing infarct size and improving cardiac function following myocardial infarction in animal models. MSC transplantation has been demonstrated to enhance angiogenesis, reduce fibrosis, and promote cardiomyocyte survival^2–9^. In experimental autoimmune encephalomyelitis (EAE) mice, MSCs modulate the immune response, reduce central nervous system inflammation, and support neurological recovery^10–15^. In rabbit and dog models of osteoarthritis, MSCs have reduced cartilage degradation, alleviated pain, and improved joint function^16–22^. As of March 2025, more than 1,800 clinical studies involving MSCs and their secretome have been registered on clinicaltrials.gov, targeting over 920 medical conditions such as Acute Respiratory Distress Syndrome (ARDS), sepsis, Graft-versus-Host Disease (GvHD), stroke, spinal cord injury, myocardial infarction, multiple sclerosis, organ transplantation, rheumatoid arthritis, Crohn’s, systemic lupus erythematosus, ulcerative colitis and COVID-19^23–35^. A meta-analysis including 55 randomized clinical studies with 2696 patients finds MSCs induce minor adverse effects^36^. Additionally, no signals of tumorgenicity and pro-coagulation risks are found^36^. Thirteen MSC-based therapies have been approved for clinical use worldwide.

However, the advancement of MSC-based therapies faces a major challenge: the difficulty of expanding large quantities of MSCs while preserving their functionality and maintaining a small cell size. Currently, MSCs are expanded as monolayers on rigid polymer substrates, including plastic flasks, polymer microcarriers in stirred tank bioreactors (STR), polymer scaffolds in packed bed (PB) bioreactors, and polymer hollow fibers in hollow fiber (HF) bioreactors^37–40^. In vivo, MSCs reside in soft 3D niches rich in cell-cell and cell-matrix interactions and autocrine and paracrine signaling^1^. Current methods create a starkly different 2D stiff microenvironment. For instance, plastic flasks have Young’s modulus of ∼100,000 kPa, far stiffer than natural soft tissues^41,42^. MSCs rapidly undergo senescence in 2D culture, losing their replicative ability and therapeutic potency^43,44^. This may explain the discrepancy between compelling preclinical data and less effective clinical outcomes^34,45–47^. While preclinical studies use young, potent MSCs, clinical trials often rely on high-passage MSCs with diminished functionality.

MSCs also enlarge during 2D culture^48–51^. A critical determinant of MSC therapeutic efficacy is their *in vivo* biodistribution. After systemic administration, MSCs frequently encounter a "first-pass effect," leading to initial entrapment in organs such as the lungs^52,53^. Cell size significantly influences this process; larger MSCs are more likely to become lodged in the microvasculature of the lung, impairing their ability to reach target tissues^51^. Moreover, oversized MSCs may cause microcirculation obstruction, ischemia, or stroke^49,54,55^. There is a pressing need for new culture strategies that can efficiently expand MSCs without causing cell enlargement and loss of function.

Recent research has found that culturing MSCs as spheroids (referred to as MSC spheroid culture) may overcome some of these limitations^56–58^. Spheroid culture better mimics the in vivo microenvironment by promoting natural extracellular matrix (ECM) composition and cell-cell interactions. It has been shown to mitigate senescence, preserve a youthful phenotype with smaller cell size, improved survival, increased secretion of trophic factors, and elevated expression of stemness-related genes^48,50,59,60^. However, as anchorage-dependent cells, MSCs typically do not proliferate in spheroid culture.

In this work, we present a novel approach that combines the benefits of both 2D and 3D spheroid culture methods to grow placenta-derived MSCs. Briefly, MSCs are expanded as adherent monolayers in 2D flasks for several days. After each passage, the cells are transitioned to a non-adherent environment for 24 to 72 hours to form 3D spheroids. For simplicity, we refer to this method as the "2D/3D culture protocol." Our hypothesis is that spheroid formation following 2D expansion can restore MSC size and function, thereby mitigating cell enlargement and senescence.

## Methods

### MSC isolation and 2D culture

Full-term human placentas were obtained from ZenBio Inc. Briefly, placentas were washed and cut into approximately 0.5 cm³ pieces, which were then partially digested with TrypLE Select solution (Gibco) at 37 °C for 30 minutes (Fig. 1A). Following digestion, 15–20 tissue pieces were transferred to a 75 cm² tissue culture flask containing 9 mL of EBM-2 complete medium (EBM-2 supplemented with 10% fetal bovine serum and 1% Penicillin-Streptomycin). The flasks were placed in an incubator and left undisturbed for three days to allow tissue attachment. After this period, the medium was replaced every three days until the cells reached confluence. These cells were designated as passage 0 (P0). P0 cells were either cryopreserved or subcultured. Details of MSC isolation and characterization can be found in our previous publication^61^.

**Figure 1.**
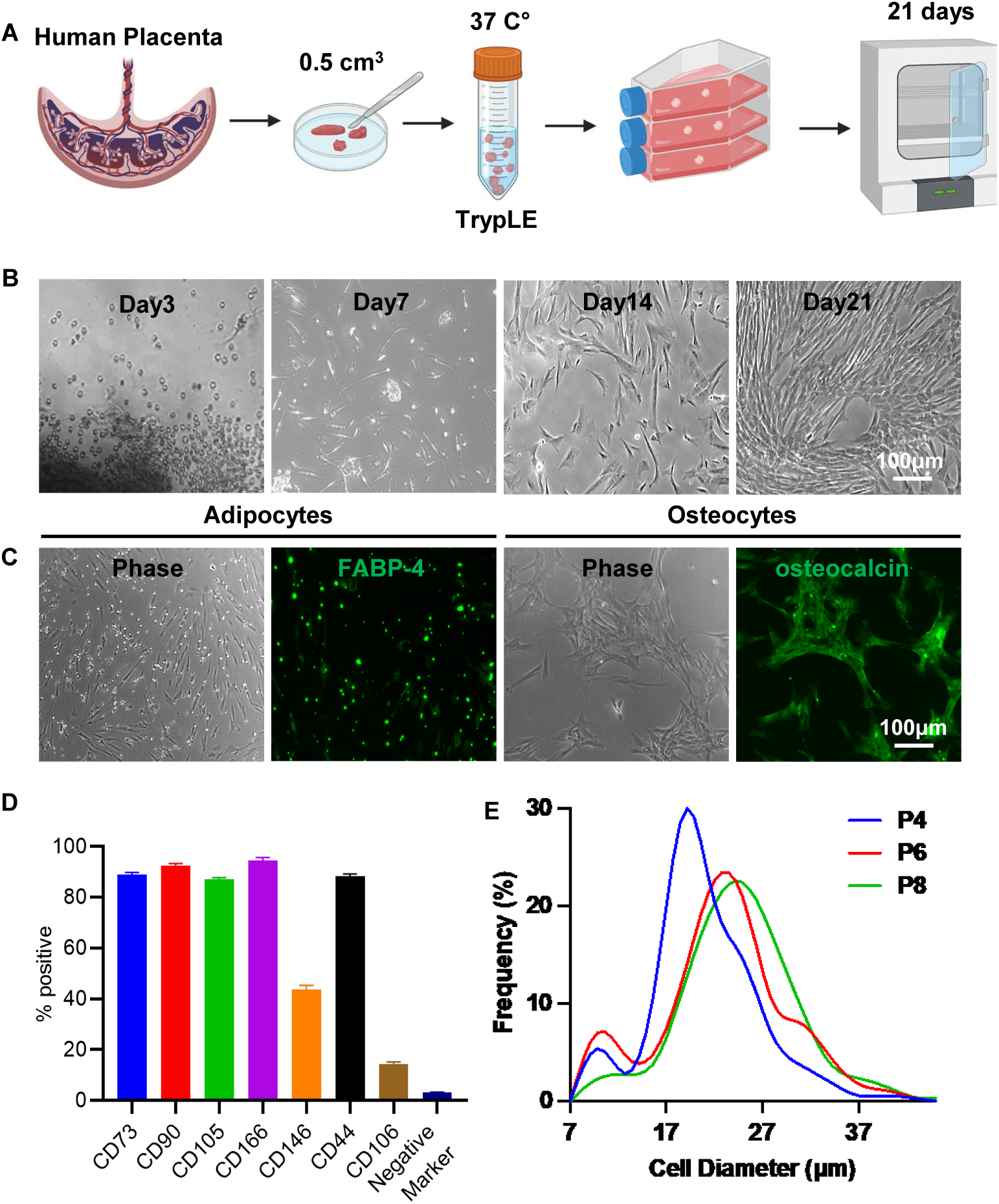
Isolation of Placenta-Derived MSCs. (A) Illustration of the MSC isolation process from human placenta. (B) Representative morphology of MSCs during isolation. (C) Differentiation of MSCs into adipocytes (FABP-4⁺) and osteocytes (osteocalcin⁺). (D) Flow cytometry analysis of MSC surface markers. Negative markers include CD34, CD45, CD11b, CD79A, and HLA-DR. (E) Size distribution of MSCs cultured in T-25 flasks at passages 4 (P4), 6 (P6), and 8 (P8).

### MSC surface marker characterization

MSCs were characterized using the Human Mesenchymal Stem Cell Verification Flow Kit (R&D Systems), which includes antibodies against the positive markers CD90, CD73, and CD105, and the negative markers CD45, CD34, CD11b, CD79A, and HLA-DR. Additionally, the Human Mesenchymal Stem Cells Multi-Color Flow Kit (R&D Systems) was used to assess expression of the positive markers CD44, CD106, CD146, and CD166. Flow cytometric analysis was performed using the BD FACSCanto™ II System.

### MSC differentiation

MSCs were evaluated for their differentiation potential using the Human Mesenchymal Stem Cell Functional Identification Kit (R&D Systems), following the manufacturer’s instructions. After 21 days of induction, cells were fixed and stained with an FABP-4 antibody to identify adipocytes and an osteocalcin antibody to identify osteocytes.

### MSC 2D culture

MSCs were seeded into T-25 flasks at a density of 8,000 cells/cm² in 4 mL of EBM-2 complete medium. The culture medium was replaced every three days. Once the cells reached approximately 90% confluency, they were harvested with 0.25% Trypsin and subcultured at the same seeding density.

### MSC 3D spheroid culture

Dissociated single MSCs were seeded into 96-well clear, round-bottom, ultra-low attachment microplates (Corning) at densities ranging from 2,000 to 40,000 cells per well in 200 μL of EBM-2 complete medium or other indicated media. The medium was refreshed every three days. To promote spheroid formation, plates were centrifuged at 100 × g for 8 minutes. Spheroids were cultured for the indicated duration. To assess cell viability, propidium iodide was added to the medium and incubated for 30 minutes prior to imaging with fluorescence microscopy.

### Alternating 2D/3D culture

MSCs harvested from 2D culture (25,000 cells per well) were seeded into 96-well clear, round-bottom, ultra-low attachment microplates to form spheroids. After 48 hours, the spheroids were dissociated into single cells, which were then seeded into 2D T-25 flasks at a density of 8,000 cells/cm² in 4 mL of EBM-2 complete medium. The medium was changed every three days. Once the cells reached approximately 90% confluency, they were harvested and cultured again as spheroids for 48 hours before being replated onto 2D flasks.

### MSC size quantification

Spheroids were harvested and washed twice with PBS. To dissociate them into single cells, 2 mL of 0.25% trypsin-EDTA (Corning) was added and incubated at 37 °C in a water bath for 10 minutes. The enzymatic reaction was stopped by adding 2 mL of EBM-2 complete medium. Cells were then centrifuged at 300 × g for 8 minutes and resuspended in EBM-2 complete medium. Images of the single cells were captured using a Zeiss fluorescence microscope. Cell size was analyzed using ImageJ software using the following procedure (Fig. S1):

- Import phase contrast images into ImageJ
- Set the image scale via Analyze → Set Scale
- Adjust threshold using Image → Adjust → Color Threshold
- Measure cell area using Analyze → Analyze Particles, setting the circularity range to 0.70–1.00
- Export the results to Excel and calculate cell diameter using the formula:

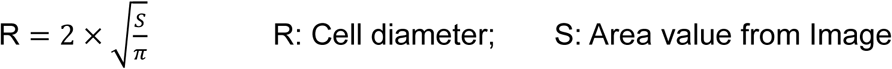

### Macrophage culture

RAW 264.7 cells (RAW-Dual™ cells, InvivoGen) were cultured in DMEM supplemented with 4.5 g/L glucose, 2 mM L-glutamine, 10% heat-inactivated fetal bovine serum (FBS), 100 µg/mL Normocin, and 1% Penicillin-Streptomycin. Cells were seeded at a density of 1.5 × 10⁴ cells/cm², and the medium was renewed twice a week.

### Macrophage inflammation assay

RAW 264.7 macrophage cells were stimulated in DMEM complete medium containing 100 ng/mL lipopolysaccharide (LPS; O111:B4, Sigma) and 10 ng/mL murine interferon-gamma (IFN-γ; Peprotech). MSCs were co-cultured with RAW 264.7 cells with the ratio as MSC/RAW: 1/10. After 24 hours, conditioned media were collected for cytokine analysis using enzyme-linked immunosorbent assays (ELISA). Additionally, the RAW-Dual™ cells is engineered to have a luciferase gene under the control of an ISG54 minimal promoter in conjunction with five IFN-stimulated response elements (ISRE). It reports the activation of interferon regulatory factors (IRFs), which contributes to inflammation. The luciferase level is measured using the commercial kit.

### Modifying Alginates with RGD peptides

A 2% (w/v) alginate solution (Cat. #194-13321, 80–120 cP, Wako Chemicals) was prepared by dissolving alginate in 0.1 N NaOH. The solution was then reacted with divinyl sulfone (DVS) at a 1:3 molar ratio of hydroxyl groups to DVS for 15 minutes. Excess DVS was removed by dialysis. Approximately 20–30% of the hydroxyl groups in the alginate polymer were successfully modified with DVS. To synthesize alginate-RGD, RGD peptides containing a C-terminal cysteine were conjugated to the DVS-modified alginate under alkaline conditions. About 10% of the modified hydroxyl groups were functionalized with RGD peptides. The resulting alginate-RGD was then blended with unmodified alginate to prepare a 2% (w/v) alginate solution, which was used to fabricate alginate hydrogel tubes.

### Processing alginate hydrogel tubes (AlgTubes)

A custom-made micro-extruder was used to process AlgTubes. A hyaluronic acid (HA) solution containing single cells and an alginate/alginate-RGD solution were pumped into the central and side channels of the micro-extruder, respectively, to form coaxial core-shell flows that were extruded into a CaCl_2_ buffer (100 mM) to form AlgTubes. If necessary, AlgTubes were further soaked in 1 mM Poly(ethylene glycol) dithiol (HS-PEG-SH, Mw3400) for 10 mins at pH=8.0 to have a secondary covalent crosslinking through the Michael addition reaction between -SH and -VS. Detailed methods for processing AlgTubes can be found in our previous publications^62–74^.

### Culturing cells in AlgTubes

For standard cell culture, 20 µL of cell-laden AlgTubes were suspended in 2 mL of culture medium in each well of a 6-well plate. Cells were seeded at a density of 1–2 × 10⁶ cells/mL within the hydrogel tube space. The tubes were formed using 2% alginate modified with 1 mM RGD peptide. The resulting hydrogel tubes had diameters ranging from 200 to 300 µm, with shell thicknesses of approximately 30–70 µm. To detach MSC from the AlgTubes, 1.2 mM free RGD peptides were added to the culture medium.

### Statistical analysis

All the data were analyzed using GraphPad Prism 8 statistical software. P value was determined by one-way analysis of variance (ANOVA) for comparison between the means of three or more groups or unpaired two-tailed t-tests for two groups analysis. The significance levels are indicated by p-value, *: p<0.05, **: p<0.01, ***: p<0.001.

## Results

### Isolation and Characterization of Placenta-Derived MSCs

MSCs were isolated from full-term placental tissue. The placenta was minced into small fragments and enzymatically digested with TrypLE for 30 minutes. The digested tissue was transferred to a T-25 cell culture flask and incubated undisturbed for 3 days to allow the fragments to adhere to the flask surface (Fig. 1A). By day 3, cells began migrating out of the tissue explants. By day 7, numerous spindle-shaped MSCs were visible, and by day 14, the cells had proliferated and spread extensively. By day 21, the culture had expanded significantly, covering approximately 70% of the flask surface (Fig. 1B). At this stage, the remaining tissue fragments were removed, and the adherent cells were expanded to full confluence. Cells were harvested using 0.25% trypsin and either cryopreserved or sub-cultured for further use.

The isolated cells exhibited the characteristic spindle-shaped morphology of MSCs and demonstrated multipotency by differentiating into FABP4-positive adipocytes and osteocalcin-positive osteocytes under appropriate conditions (Fig. 1C). Flow cytometry analysis of cells showed that most of them expressed classical MSC surface markers, including CD73, CD90, CD105, CD44, and CD166, while expression of hematopoietic and immune markers—such as CD45, CD34, CD11b, CD79A, and HLA-DR—was negligible (Fig. 1D). In our previous studies^61^, we also demonstrated that these cells possess strong immunomodulatory capabilities: they suppressed cytokine release from macrophages, inhibited reactive oxygen species (ROS) production and neutrophil extracellular trap (NET) formation by neutrophils, and reduced T cell proliferation under inflammatory conditions in vitro. In an acute lung injury mouse model, these MSCs significantly lowered cytokine levels, reduced tissue damage, and improved survival outcomes.

Furthermore, these MSCs were cultured over multiple passages, and cell size (diameter) distribution was quantified after each harvest. A clear trend of increasing cell size with successive passages was observed, indicating progressive cellular enlargement with continued replication (Fig. 1E). This observation is consistent with previous reports that MSCs tend to enlarge during in vitro culture on stiff 2D surfaces. In short, we have isolated authentic MSCs from placenta.

### Spheroid Culture Reduce Placenta MSC Size

We first investigated whether placenta MSCs could form spheroids. MSCs were seeded into low attachment 96-well plates at densities of 2,000, 10,000, 25,000, and 40,000 cells per well. All groups successfully formed spheroids within 24 hours (Fig. 2A). At this time point, the average diameters of the spheroids were as follows: 191.0 µm for 2K, 338.1 µm for 10K, 443.1 µm for 25K, and 462.7 µm for 40K. Notably, the 2K spheroids maintained a consistent diameter over extended culture periods (48 and 72 hours), whereas the diameters of the 10K, 25K, and 40K spheroids decreased with prolonged culture (Fig. 2B).

**Figure 2.**
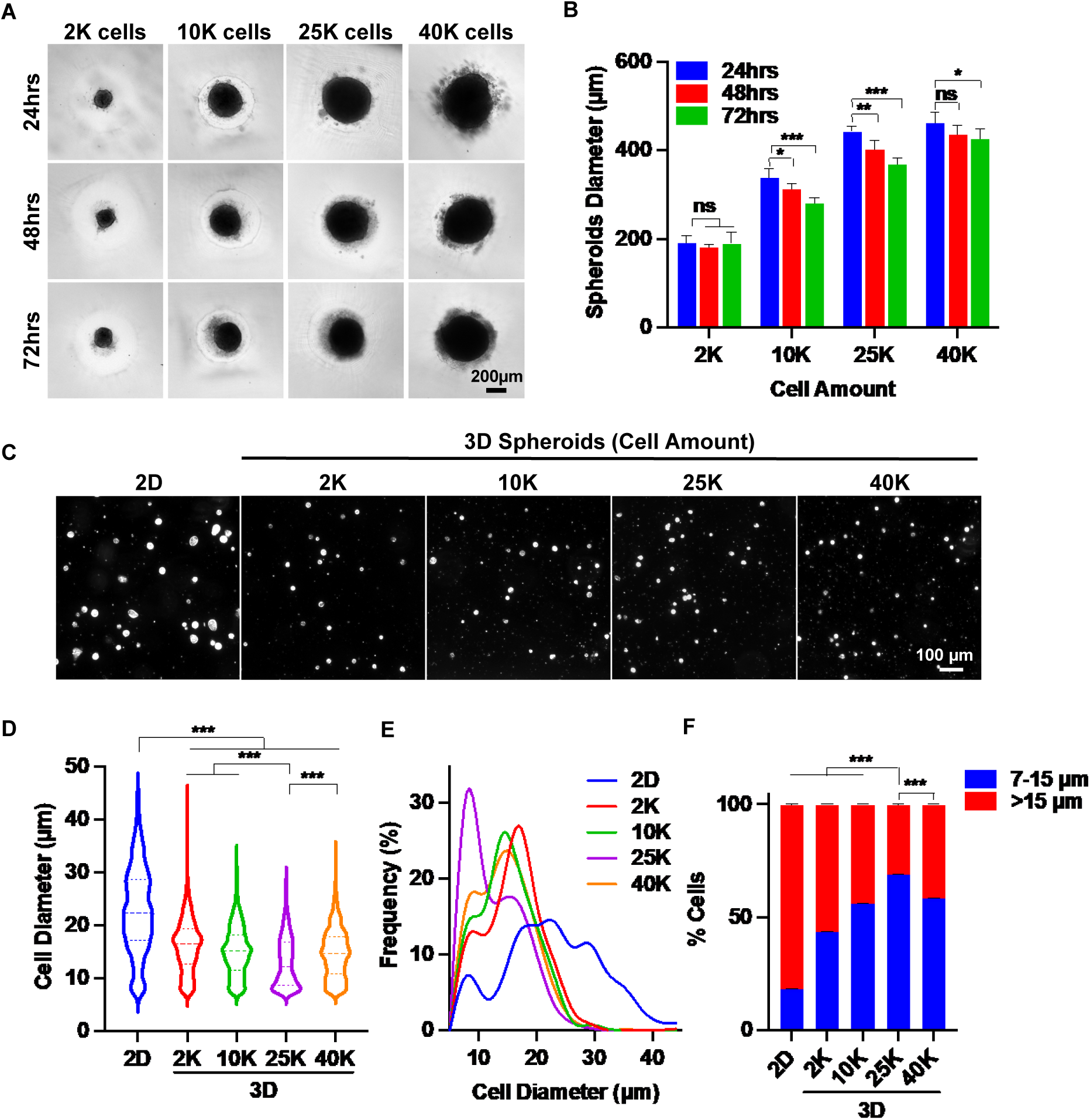
3D Spheroid Culture Reduces MSC Size: Identifying the Optimal Spheroid Diameter. (A) Representative images of MSC (P3) spheroids cultured for 3 days with varying initial cell numbers per spheroid. (B) Quantification of spheroid diameter over the 3-day culture period. (C) Representative images of individual MSCs following 2D and 3D spheroid culture. (D, E) Cell size distribution of MSCs after 2D and 3D spheroid culture. (F) Proportions of small (≤15 µm) and large (>15 µm) MSCs after 2D and 3D spheroid culture. Spheroid culture duration in panels (C–F) was 72 hours.

Next, we assessed whether spheroid culture reduced the size of individual MSCs. Spheroids were dissociated into single cells and imaged using phase-contrast microscopy. Regardless of the spheroid size, MSCs derived from 3D spheroid cultures appeared markedly smaller than those from 2D cultures (Fig. 2C). Using ImageJ software, we measured the diameters of over 600 MSCs per group, following a standardized protocol (Fig. S1). Cell diameter distributions were visualized using Prism 9 software. The mean diameter of MSCs from 2D culture was 22.8 µm. In contrast, MSCs from 2K, 10K, 25K, and 40K spheroids cultured for 72 hours had mean diameters of 16.1 µm, 15.1 µm, 13.0 µm, and 14.6 µm, respectively—significantly smaller than those from 2D culture (Fig. 2D). Additionally, MSCs from 2D culture exhibited a much broader size distribution compared to those from spheroid cultures.

Interestingly, size distribution histograms revealed two distinct peaks (Fig. 2E). The proportion of cells in the smaller size peak was significantly higher in 3D spheroid cultures than in 2D cultures. We categorized MSCs based on diameter: cells between 7–15 µm were defined as "small MSCs," while those larger than 15 µm were classified as "big MSCs." In 2D cultures, small MSCs comprised less than 20% of the population, with big MSCs accounting for over 80% (Fig. 2F). In contrast, 3D cultures showed a marked increase in the proportion of small MSCs, reaching at least 40%, while big MSCs dropped below 60%. Among the 3D groups, the 25K spheroids yielded the highest proportion of small MSCs and the lowest proportion of big MSCs, indicating that this condition was most effective in reducing cell size. Therefore, 25K spheroids were used in subsequent experiments.

We then systematically examined the effect of spheroid culture duration on MSC size. Spheroids formed from 25K cells were cultured for up to seven days. While spheroid diameter gradually decreased over time, structural integrity was maintained throughout the 7-day period (Fig. 3A, C). Spheroids were dissociated daily to measure changes in cell size (Fig. 3B). In contrast to 2D cultures, where MSC size steadily increased, spheroid-cultured MSCs exhibited a significant size reduction within the first 48 hours. Although the size continued to decrease from 48 to 168 hours, the rate of reduction was less pronounced (Fig. 3D). Consistent with the reduction in mean diameter, the size distribution of MSCs became progressively narrower with longer culture durations (Fig. 3E, F). These findings suggest that a spheroid culture duration of 48 to 72 hours is optimal for reducing MSC size.

**Figure 3.**
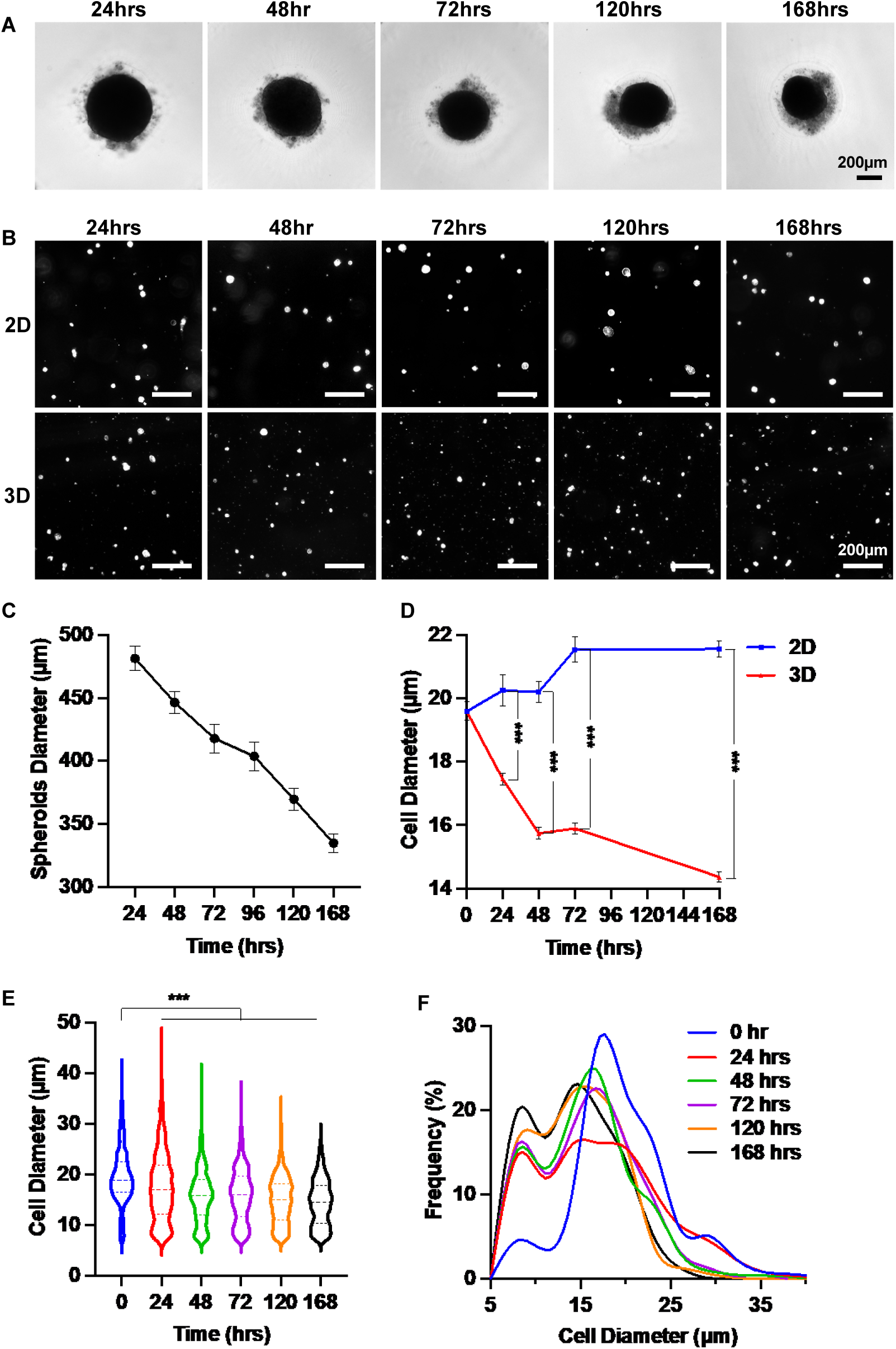
3D Spheroid Culture Reduces MSC Size: Determining the Optimal Culture Duration. (A) Representative images showing morphological changes in MSC (P4) spheroids over a 7-day culture period. (B) Images of individual MSCs cultured in 2D or in 25K-cell spheroids for varying durations. (C) Changes in the diameter of 25K-cell MSC 3D spheroids over 7 days. (D) Comparison of mean MSC diameter in 2D versus 25K spheroid cultures over 7 days. (E, F) Size distribution of MSCs within 25K spheroids across the 7-day culture period.

### Spheroid Culture Can Be Performed in Chemically Defined Medium

During spheroid culture, we observed that not all MSCs were incorporated into the spheroids. Some cells remained loosely attached to the spheroid surface. To assess their viability, we stained the spheroids with Propidium Iodide (PI), which labels dead cells with red fluorescence. These loosely associated cells were confirmed to be non-viable (Fig. 4A). We hypothesized that supplementing the culture with extracellular matrix (ECM) proteins might enhance spheroid formation and cell viability. MSCs were seeded at 25,000 cells per well and cultured in EBM-2 complete medium, with or without 0.625 µg/mL iMatrix-511 (Nippi), 0.625 µg/mL iMatrix-411 (Nippi), 0.625 µg/mL rhLaminin-521(AMSBIO), and 6.25 µg/mL fibronectin (Sigma, F1141). Our results demonstrated that ECM proteins significantly improved spheroid formation and reduced the presence of dead cells (Fig. 4A). In the absence of ECM proteins, approximately 30% of spheroids exhibited dead cell aggregates, whereas this number dropped to 10% with ECM supplementation (Fig. 4B). Furthermore, ECM proteins reduced the diameter of individual MSCs (Fig. 4C) and significantly increased the proportion of small MSCs (Fig. 4D).

**Figure 4.**
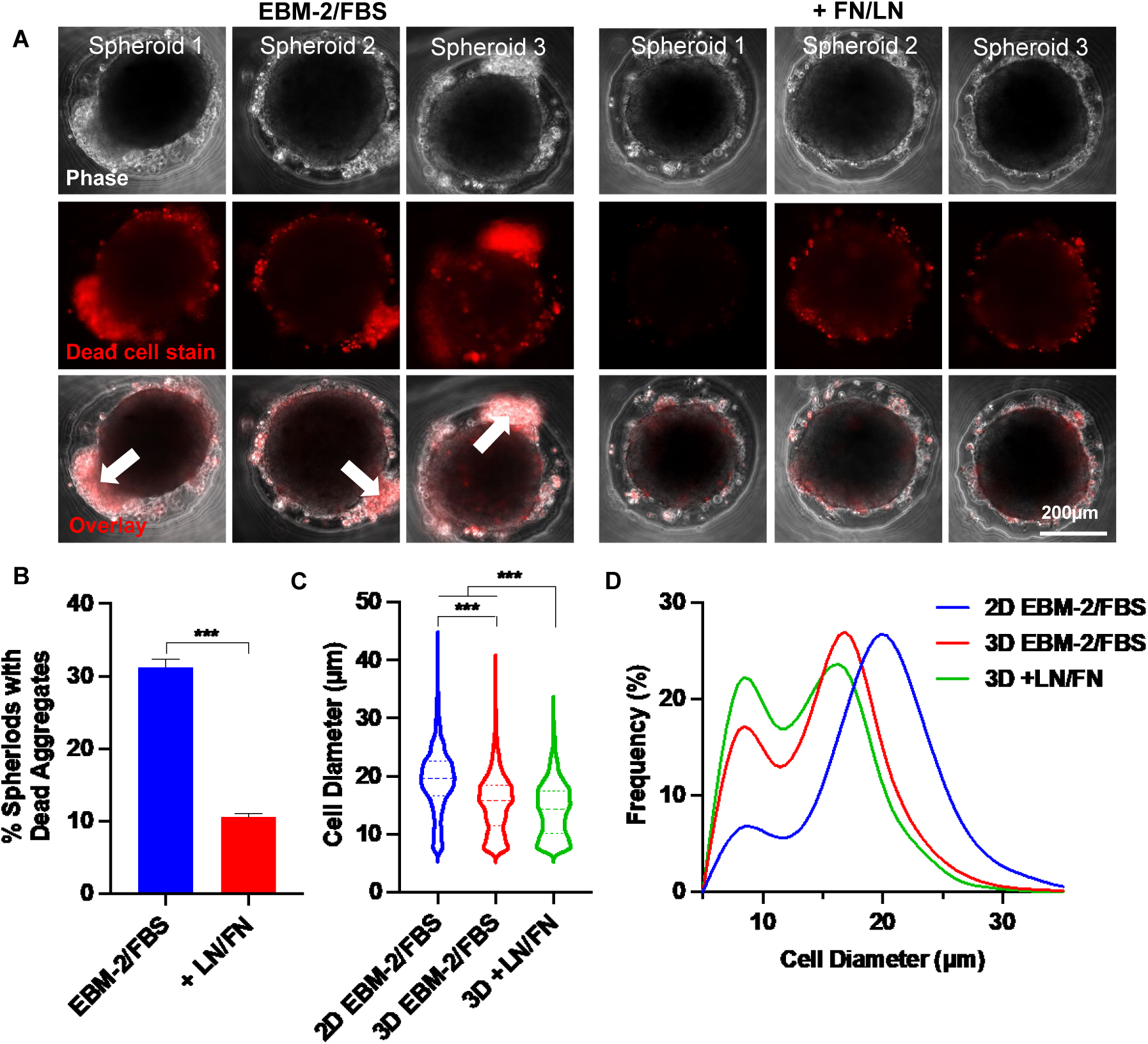
Effects of Supplementing Extracellular Matrix Proteins on MSC Viability and Size in Spheroid Culture. (A) Phase-contrast and dead cell staining (red) images of MSC (P4) spheroids cultured in EBM-2 + 10% FBS medium, with or without laminin (LN) and fibronectin (FN). Three representative spheroids are shown in each condition. The white arrow indicates the loosely associated cells. (B) Percentage of spheroids containing dead cell aggregates in the presence or absence of LN and FN. (C, D) MSC size distribution following 2D monolayer and 3D spheroid cultures. Spheroids were cultured for 48 hours.

The medium used in the above experiments contained 10% fetal bovine serum (FBS). To determine whether spheroids could be formed in a chemically defined medium, MSCs were seeded at 25,000 cells per well and cultured in three different media: EBM-2 + 10% FBS (EBM-2 + FBS), EBM-2 basal medium supplemented with insulin-transferrin-selenium (ITS) (EMB-2 + ITS) and DMEM/F12 supplemented with ITS (DMEM/F12 + ITS). Our data showed a high number of dead cells in the FBS-containing medium (Fig. 5A). Replacing FBS with ITS did not impair spheroid formation and significantly reduced cell death. Remarkably, spheroids cultured in DMEM/F12 + ITS exhibited almost no dead cells. Despite these differences in viability, the diameters of MSCs across the three media were comparable (Fig. 5B, C).

**Figure 5.**
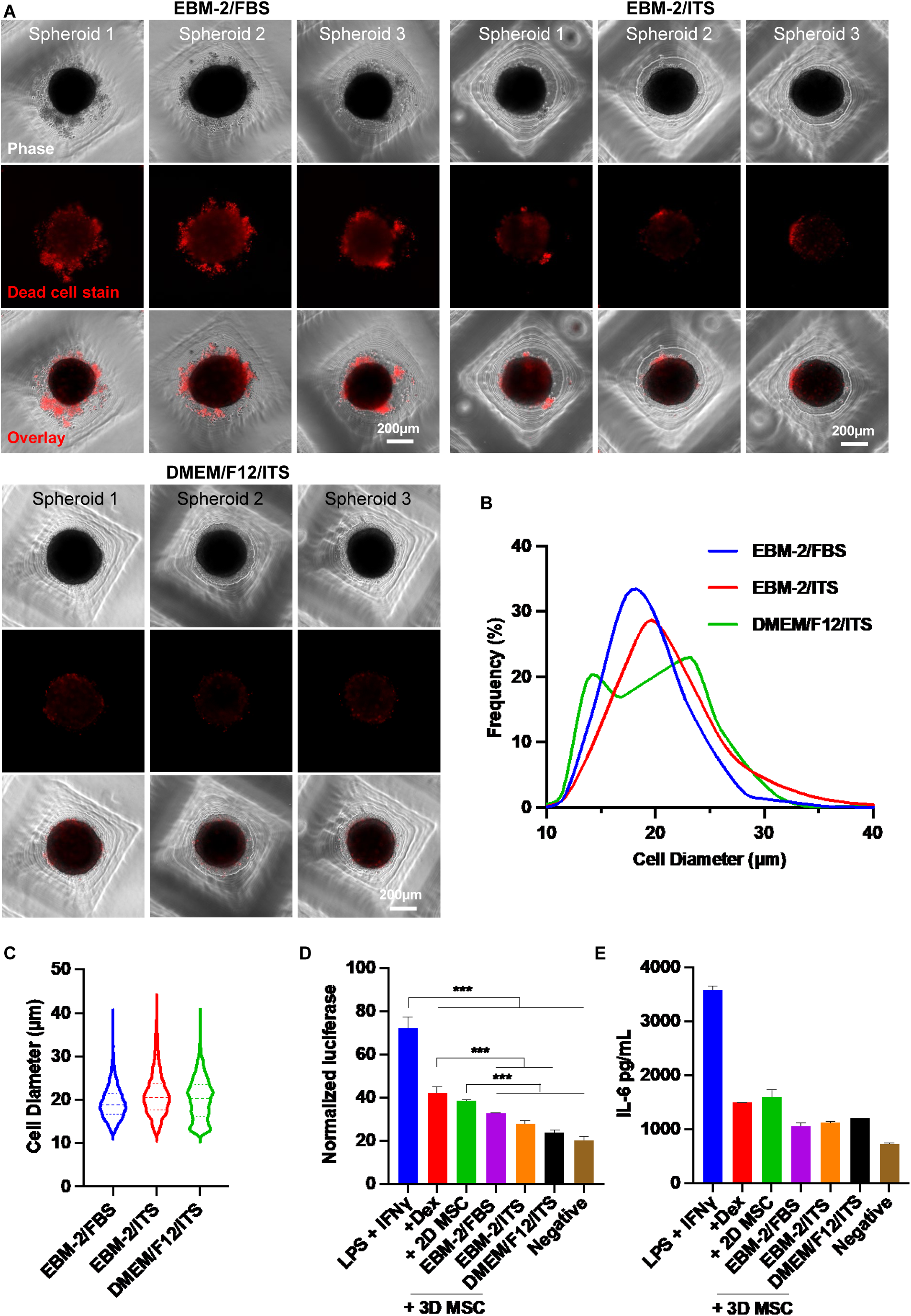
Effect of Chemically Defined Medium on MSC Viability and Size in Spheroid Culture. (A) Phase-contrast and dead cell staining (red) images of MSC (P6) spheroids cultured in various media. Three representative spheroids are shown in each condition. (B, C) MSC size distribution in 3D cultures across different media conditions. (D, E) Anti-inflammatory capability of MSCs cultured under varying conditions. RAW-Dual reporter cells were stimulated with 100 ng/mL LPS and 10 ng/mL IFNγ, then treated with MSCs. Dexamethasone (Dex, 1 µg/mL) was used as a positive control. Luciferase activity (D) and mouse IL-6 (E) were measured. Spheroid culture time was 48 hrs.

To evaluate whether the defined medium affected MSC function, we assessed their ability to suppress macrophage-mediated inflammation (Fig. 5D, E). We focused on the interferon regulatory factor (IRF) signaling pathway, a key regulator of inflammation. RAW 264.7 macrophages engineered to express a secreted luciferase reporter under IRF control were used. Cells were stimulated with 100 ng/mL LPS and 10 ng/mL IFNγ to induce inflammation. Inflamed macrophages were then treated with MSCs. Dexamethasone (1 µg/mL), a clinically used anti-inflammatory agent, served as a benchmark. After 24 hours, the luciferase activity in medium was quantified. The pro-inflammatory cytokine IL-6 was also measured using ELISA, with antibodies specific to mouse IL-6 to avoid cross-reactivity with human cytokines secreted by MSCs. All treatments—including MSCs from both 2D and 3D cultures—reduced IL-6 and luciferase expression (Fig. 5D, E). MSCs from 2D culture showed similar efficacy to dexamethasone, while MSCs from spheroid culture outperformed dexamethasone. Importantly, no significant differences were observed among the three media conditions, indicating that chemically defined media do not compromise MSC immunomodulatory function.

### Alternating 2D Flask and 3D Spheroid Culture Slows MSC Size Enlargement and Senescence

Previous results demonstrated that spheroid culture reduces MSC size and enhances their functional properties. However, because MSCs are anchorage-dependent, they do not proliferate in non-adherent spheroid conditions. To address this limitation, we tested a new culture protocol that alternates between 2D and 3D environments—leveraging the proliferative capacity of 2D culture and the size-reducing, function-restoring benefits of 3D spheroid culture. In this protocol, MSCs are first expanded in standard 2D flasks. After each passage, cells are transferred to ultra-low attachment plates to form spheroids for 48 hours. Although MSCs do not proliferate during the spheroid phase, their size is significantly reduced (Fig. 6A).

**Figure 6.**
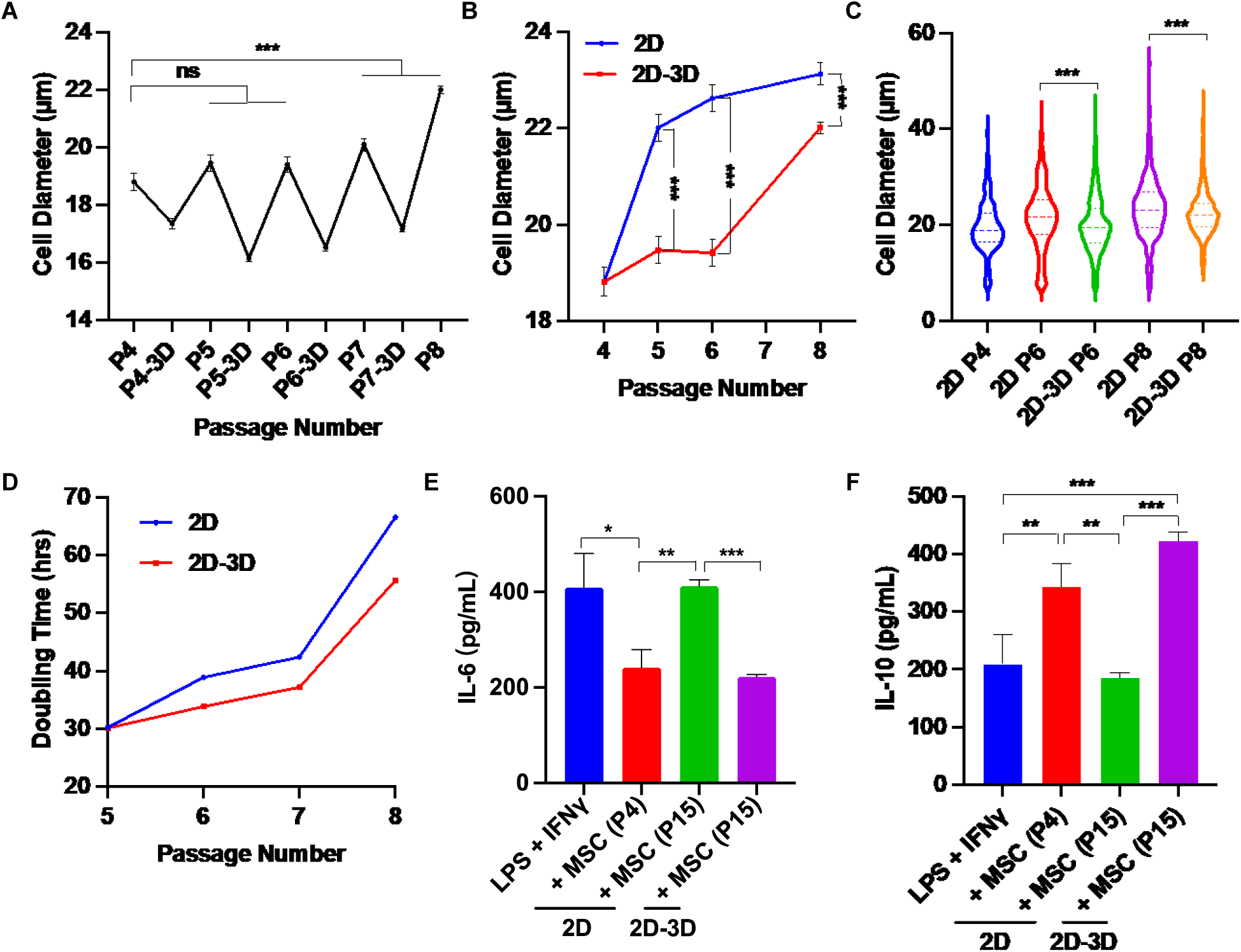
Effects of Alternating 2D/3D Culture on MSC Size and Immunomodulatory Function. (A) MSCs were cultured in flasks for four passages, with an additional 2-day spheroid culture following each passage. Shown are the mean cell diameters immediately after 2D culture and after subsequent spheroid culture. (B) MSC diameters from P5 to P8 using conventional 2D culture versus the alternating 2D/3D method. P4 cells served as the starting population for both conditions. Diameters were measured after harvesting from 2D flasks. (C) Comparison of MSC sizes at P5–P8 between the two culture methods. (D) Comparison of doubling times at P5–P8 between the two methods. (E, F) Macrophages were activated with LPS and IFNγ, then treated with either early-passage (P4) or late-passage (P15) MSCs derived from conventional 2D culture or the alternating 2D/3D protocol. Levels of mouse IL-6 and IL-10 were measured to assess immunomodulatory effects.

We compared long-term culture outcomes between continuous 2D culture and the alternating 2D/3D protocol. In 2D culture, MSC diameter increased significantly by passage 5. In contrast, MSCs cultured using the 2D/3D protocol maintained a smaller size until passage 8 (Fig. 6B, C). At every passage, MSCs from the 2D/3D protocol were consistently smaller than those from 2D culture alone. Additionally, the 2D/3D protocol resulted in shorter population doubling times, indicating improved proliferative efficiency.

To assess the impact of long-term culture on MSC immunomodulatory function, cells were expanded to passage 15 (P15) using both methods. MSCs from 2D passage 4 (2D P4), 2D P15, and 2D/3D P15 were co-cultured with RAW 264.7 macrophages, and cytokine levels were measured by ELISA. For IL-6, a pro-inflammatory cytokine, secretion levels in the 2D P4 and 2D/3D P15 groups were comparable and significantly lower than those in the positive control group. In contrast, IL-6 levels in the 2D P15 group were similar to the positive control, indicating a loss of immunosuppressive function (Fig. 6E). For IL-10, an anti-inflammatory cytokine, 2D P15 MSCs secreted levels comparable to the positive control, suggesting diminished function. Notably, 2D/3D P15 MSCs secreted higher levels of IL-10 than even the 2D P4 group, indicating preserved or enhanced immunomodulatory capacity (Fig. 6F). These findings suggest that alternating 2D and 3D culture phases can effectively delay MSC enlargement and senescence while preserving their therapeutic potential.

## Discussion

### Mesenchymal Stem Cells Undergo Enlargement and Senescence in 2D Culture

MSCs are widely used in tissue engineering, regenerative medicine, and cell-based therapies. To meet the demand for large cell quantities, MSCs are typically expanded in vitro using 2D culture flasks, where they adhere and proliferate on plastic surfaces^75^. However, prolonged culture in these artificial environments often leads to progressive cell enlargement, reduced proliferation, and functional decline^76^. Understanding the mechanisms behind these changes is essential for preserving MSC quality and therapeutic efficacy^75^.

A primary contributor to MSC enlargement is cellular senescence. Like other somatic cells, MSCs have a finite replicative lifespan and eventually enter senescence after repeated divisions^76^. This process is characterized by a shift from the typical spindle-shaped morphology to an enlarged, flattened appearance^76^. Senescent MSCs exhibit increased senescence-associated β-galactosidase (SA-β-gal) activity and a higher proportion of cells arrested in the G0/G1 phase^76^. This cell cycle arrest, combined with continued biosynthetic activity, contributes to increased cell size.

Senescence is driven by multiple interconnected processes^77–80^. These include DNA damage accumulation, telomere shortening, mitochondrial dysfunction, impaired autophagy, and epigenetic alterations^77–80^. DNA damage activates the DNA damage response (DDR) and downstream pathways such as p53/p21 and p16/Rb, leading to cell cycle arrest^77–80^. Similarly, telomere attrition from repeated replication triggers DDR and senescence^77–80^. Mitochondrial dysfunction and defective autophagy result in the buildup of damaged cellular components, further promoting senescence and morphological changes^77–80^. Oxidative stress also plays a central role by inducing DNA damage and altering cell behavior^76–80^.

In addition to intrinsic aging processes, the non-physiological conditions of 2D culture contribute to MSC enlargement. The rigid tissue culture plastic (TCP) used in flasks lacks the 3D architecture and mechanical cues of the native MSC niche^56^. In vivo, MSCs reside in a soft, ECM-rich environment that supports dynamic cell-cell and cell-matrix interactions^81^. In contrast, 2D culture exposes cells to a flat, stiff surface, which alters morphology, gene expression, and function^81^. The stiffness of TCP (in the gigapascal range) far exceeds that of native tissues (typically <0.1 kPa in 3D spheroids)^56^, promoting excessive cell spreading and intracellular tension^56^. Moreover, the lack of vertical support in 2D culture induces unnatural apical-basal polarity, which can disrupt signaling and further influence cell shape and size^82^.

However, several recent well-designed studies challenge this conventional view and suggest that cell size enlargement may actually lead to cellular senescence^83,84^. Human cells tend to grow slightly larger over their lifespans because, during each division, they pause to check for DNA damage. If damage is detected, the cell halts division to make repairs. These delays allow the cell to grow incrementally larger. Over time, with repeated divisions and repair pauses, cells accumulate size. As cells enlarge, their DNA and protein synthesis machinery struggle to meet the demands of the increased volume. This imbalance leads to insufficient protein production, resulting in cytoplasmic dilution and disruption of normal cell division. Diluted cytoplasm slows down biochemical reaction rates, as lower concentrations of key proteins may prevent certain reactions from occurring altogether. Based on this emerging theory, strategies that reduce cell size may help delay the onset of senescence.

### 3D Spheroid Culture Reduces Cell Size and Prolongs Lifespan

Recent studies suggest that 3D spheroid culture may help overcome limitations of traditional 2D MSC expansion^58,85,86^. In this system, MSCs self-assemble into spherical aggregates, promoting enhanced cell-cell and cell-matrix interactions that better mimic the native tissue microenvironment^58,85,86^. A key observation is the significant reduction in MSC size compared to cells grown in 2D monolayers^87^. This reduction in cell size is not merely morphological—it may enhance therapeutic potential. Evidence links smaller MSCs in spheroids to improved functional properties, including increased secretion of therapeutic factors and better survival after transplantation. The size reduction observed in 3D spheroids is driven by multiple factors: cytoskeletal reorganization^87^, a shift from cell-ECM to cell-cell adhesion^87^, the softer mechanical environment of spheroids^87^, and metabolic constraints due to limited nutrient diffusion^87^. A recent study proposes that the size decrease is due to increased excretion of extracellular vesicles^48^. They found that 3D culture led to more vesicles both on the cell surface and in the surrounding medium. This increased shedding appears linked to lower levels of F-actin, a component of the cell’s internal skeleton, suggesting that reducing internal tension facilitates vesicle release and subsequent cell size reduction. Treatments that also reduced F-actin levels resulted in increased vesicle secretion and smaller cells, supporting this mechanism.

### Alternating Two- and Three-Dimensional Culture Technique

However, spheroid culture alone is insufficient for large-scale MSC expansion, as MSCs proliferate poorly in this format^87^. To address this limitation, a recent study proposed a cyclical aggregation strategy that alternates between 3D aggregation and 2D expansion^60^. This approach maintains MSC viability and functionality over extended periods, preserving essential stem cell characteristics and differentiation potential. The study also identified the integrated stress response (ISR) pathway as a key mechanism underlying the beneficial effects of aggregation. However, the study focused exclusively on adipose-derived MSCs, and its applicability to MSCs from other tissue sources remains unknown. Additionally, it did not assess whether cyclical aggregation reduces cell size or enhances functional outcomes.

Among various tissue sources, the placenta is particularly attractive for MSC isolation due to its abundance and the large number of cells that can be harvested from a single placenta. Over the past decade, our team has focused on using placenta-derived MSCs to treat a range of conditions, with a particular emphasis on spina bifida. Myelomeningocele (MMC), or spina bifida, is caused by incomplete neural tube closure during spinal cord development^88–104^. Intrauterine damage to the exposed spinal cord leads to lifelong paralysis, incontinence, musculoskeletal deformities, and cognitive impairments. Our in vitro studies demonstrated that placenta MSCs protected neurons from various insults^90^. Using a retinoic acid (RA)-induced fetal rat MMC model, we found that in utero treatment with placenta MSCs significantly reduced spinal cord compression and neuronal apoptosis^97^. We validated these results in a fetal lamb model of MMC ^88–93,98,100,102,104^. Based on these findings, the Phase 1 Cellular Therapy for In Utero Repair of Myelomeningocele (CuRe) Trial was started in spring 2021 with seven patients. The FDA and the Data Monitoring Board subsequently determined that the therapy is safe. In 2023, a Phase 2 trial was launched to enroll 28 more patients to assess efficacy.

Throughout our journey with the CuRe trial, we have encountered all the MSC culture challenges. While we are currently able to produce sufficient low-passage, functional MSCs using 2D flasks for this proof-of-concept Phase 1/2a trial - involving just 35 patients and a modest dose of 1×10⁶ MSCs per patient –existing technologies are clearly inadequate to meet the demands of broader clinical applications following FDA approval. This limitation becomes even more critical when considering treatment for prevalent conditions such as stroke or myocardial infarction, which affect millions and require substantially higher cell doses (∼10⁹ MSCs per patient). There is an urgent need for transformative MSC culture technologies to unlock the full therapeutic potential of MSCs.

The alternating 2D/3D culture technique shows promise in addressing current limitations in MSC expansion. In this study, we demonstrate that placenta-derived MSCs can successfully form spheroids, and that spheroid culture significantly reduces cell size. Both spheroid size and culture duration influence MSC size (Fig. 2, 3), consistent with findings reported in the literature. Moreover, supplementing the culture with ECM proteins such as laminin and fibronectin further reduces cell death and enhances the reduction in cell size (Fig. 4). We also show that spheroid culture can be performed under fully chemically defined conditions, resulting in minimal cell death while preserving MSC function (Fig. 5). Importantly, placenta MSCs cultured using the alternating 2D/3D approach exhibit slower progression of cell enlargement and maintain proliferative capacity, in contrast to the decline typically seen in prolonged 2D culture. These MSCs also retain their immunomodulatory function, whereas their 2D-cultured counterparts lose this capability entirely (Fig. 6).

### A Potential Scalable Strategy for Leveraging the Alternating 2D/3D Culture Approach

While our results show that the alternating 2D/3D approach can significantly improve the MSC culture, the protocol is hard to be implemented at large scales. Adding a 3D spheroid culture significantly increases the process complexity and cost. New culture systems that allow both 2D and 3D culture would be helpful to implement the new approach. This need might be met with the hydrogel tube microbioreactors recently developed by our lab^62,63,72–74,64–71^. This method cultivates cells in hollow, microscale alginate hydrogel tubes. AlgTubes provide a cell-friendly microenvironment, resulting in paradigm-shifting improvements in cell viability, growth rate, yield, culture consistency, and scalability. When culturing human stem cells, we have achieved up to 4000-fold expansion per passage and 5x10^8^ cells/mL volumetric yield, which is ∼250 times the current state-of-the-art. However, AlgTubes do not support the growth of anchor-dependent stem cells, as they lack adhesion points.

To address the problem, we have developed chemistry to functionalize AlgTubes with RGD peptides that can bind integrin receptors in preliminary studies. Briefly, alginates are reacted with Divinyl Sulfone (DVS). About 30% of the OH groups are conjugated with vinyl sulfones (-VS) (Fig. 7A). RGD peptides with a thiol (-SH) group at the C-terminal are then reacted with alginate-VS to prepare RGD-modified alginates (RGD-alginate). The RGD/VS ratio controls the degree of RGD modification. Alginate-RGD and unmodified alginate are mixed and extruded into a CaCl₂ buffer, where they are rapidly crosslinked to form hydrogel tubes through ionic interactions between Ca²⁺ ions and the carboxyl groups on the alginate chains (Fig. 7B). The final RGD concentration in the AlgTubes is controlled by the proportion of alginate-RGD used in the mixture. The AlgTubes are then soaked in poly(ethylene glycol) dithiol (HS-PEG-SH) to achieve secondary covalent crosslinking via a Michael addition reaction between –SH and –VS groups (Fig. 7C). This chemistry is simple, cost-effective, biocompatible, and scalable.

**Figure 7.**
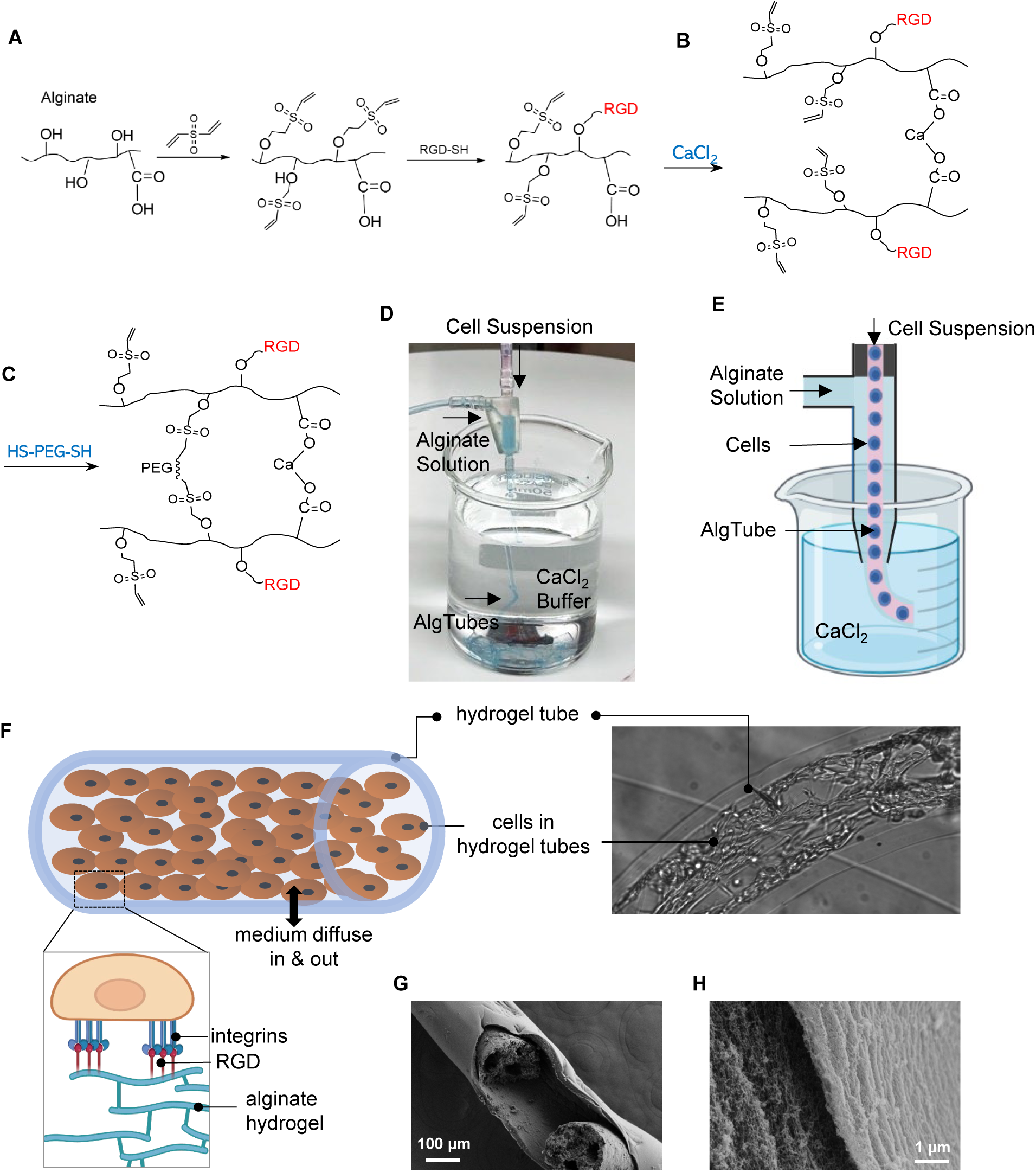
RGD-Modified Alginate Hydrogel Tubes (AlgTubes) for MSC Culture. (A) Schematic of alginate modification with RGD peptides. (B, C) Chemistry underlying hydrogel tube formation. (D, E) Process AlgTubes using a microextruder: a cell suspension and alginate solution are pumped into the central and side channels, respectively, creating coaxial core-shell flows. These are extruded through a nozzle into a CaCl₂ buffer, where Ca²⁺ ions crosslink the outer alginate shell, forming hydrogel tubes instantly. (F) Illustration of growing MSCs within an AlgTube. (G, H) SEM images showing the porous structure of the AlgTubes.

We have designed a micro-extruder to fabricate alginate hydrogel tubes and load cells into them (Fig. 7D, E). The incorporation of RGD peptides enables cells to adhere to the inner surface of the tubes, while the hydrogel provides a soft substrate for cell attachment (Fig. 7F). These hydrogel tubes contain abundant nanopores that permit the diffusion of nutrients and growth factors with molecular weights below 1000 kDa, supporting cell viability and growth (Fig. 7G, H). We developed a method to mimic the alternating 2D/3D culture protocol. MSCs were initially cultured in RGD-functionalized AlgTubes, forming a monolayer along the inner surface (Fig. 8A–C), corresponding to the 2D culture phase. Free RGD peptides were then added to the medium to competitively inhibit integrin receptors, prompting MSCs to contract within 24 hours and form spheroids by 48 hours (Fig. 8D–F), representing the 3D spheroid culture phase. Upon removal of the free RGD peptides, MSCs reattached and spread, forming a multilayered cell mass within 48 hours (Fig. 8G, H). Thus, our AlgTube system supports both adherent and suspension MSC culture modes. Future work should focus on systematically optimizing this system and characterizing MSC behavior and function within it.

**Figure 8.**
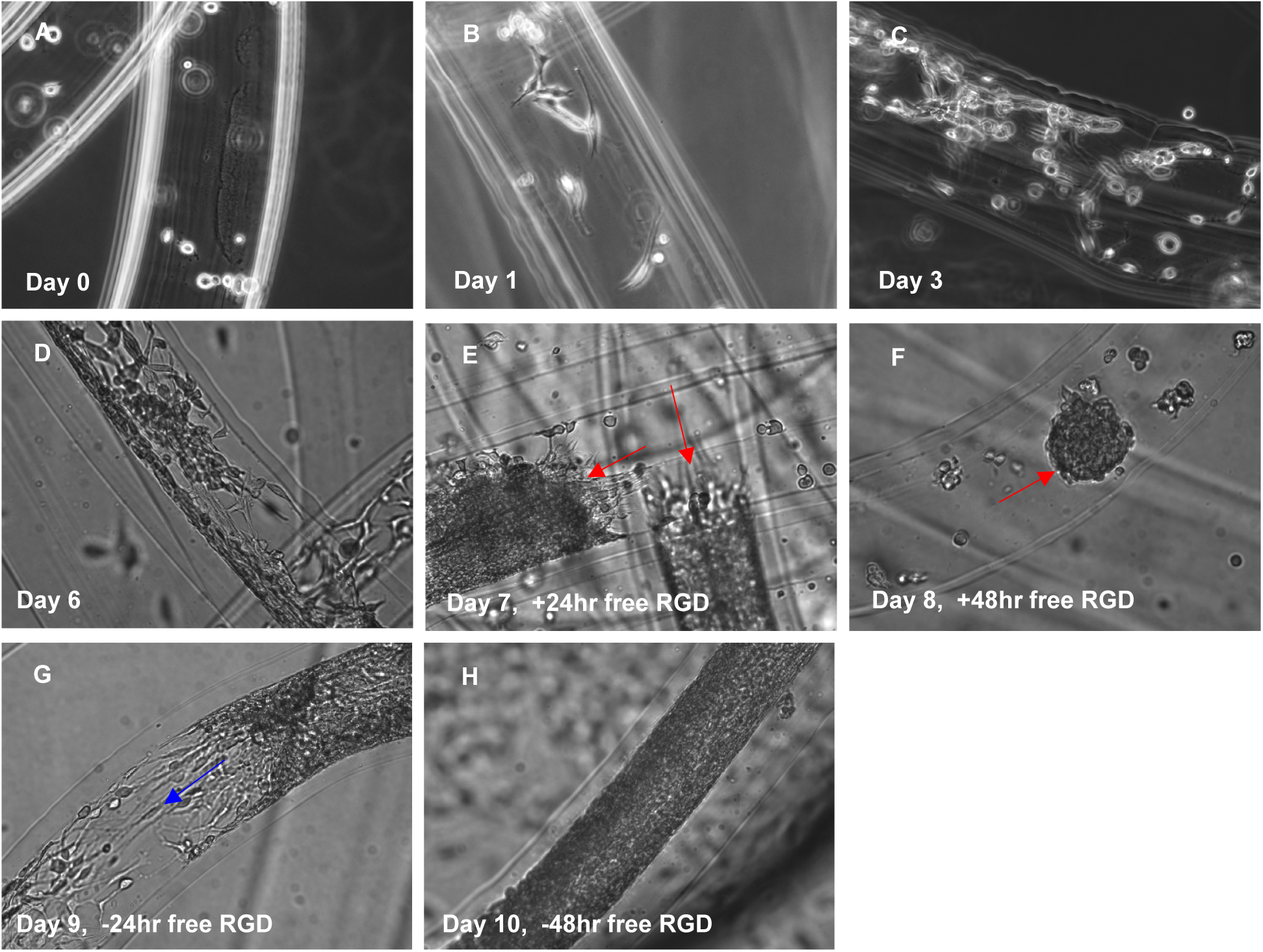
Dynamic Cell Adhesion in RGD-Modified AlgTubes. (A) MSCs were processed into RGD-modified AlgTubes. (B–D) Cells adhered to the inner surface and proliferated from day 0 to day 6. (E) On day 6, free RGD peptides were added to the culture medium, leading to cell contraction within 24 hours. (F) By 48 hours, cells had fully detached and formed spheroids (red arrows). (G, H) On day 8, removal of free RGD peptides resulted in MSC reattachment to the AlgTube surface and resumed cell growth (blue arrows).

## Conclusion

This study demonstrates that alternating 2D and 3D spheroid culture effectively mitigates placenta MSC enlargement and senescence, preserving their proliferative and immunomodulatory functions. MSCs cultured using this approach maintain a smaller size and enhanced therapeutic potential compared to conventional 2D-expanded cells. The protocol is compatible with chemically defined media and can be further optimized with ECM supplementation. To address scalability, we introduced a novel RGD-functionalized AlgTube system that supports dynamic adhesion and mimics the 2D/3D protocol, offering a versatile and scalable platform for MSC expansion. Together, these findings provide a foundation for next-generation MSC manufacturing strategies that can meet the demands of large-scale clinical applications.

## Author Contributions

Y.L. and S.W. conceived the idea. Y.L., S.W., Y.P., L.H., A.W., L.X., W.Z. designed the experiments. Y.P., L.H., Y.Y., X.W. conducted experiments. Y.L., S.W., Y.P., L.H., A.W., L.X., W.Z., W.L. carried out data analysis and manuscript preparation.

## Data Availability

The authors confirm that the data supporting the findings of this study are available within the article.

---

**Figure S1.**
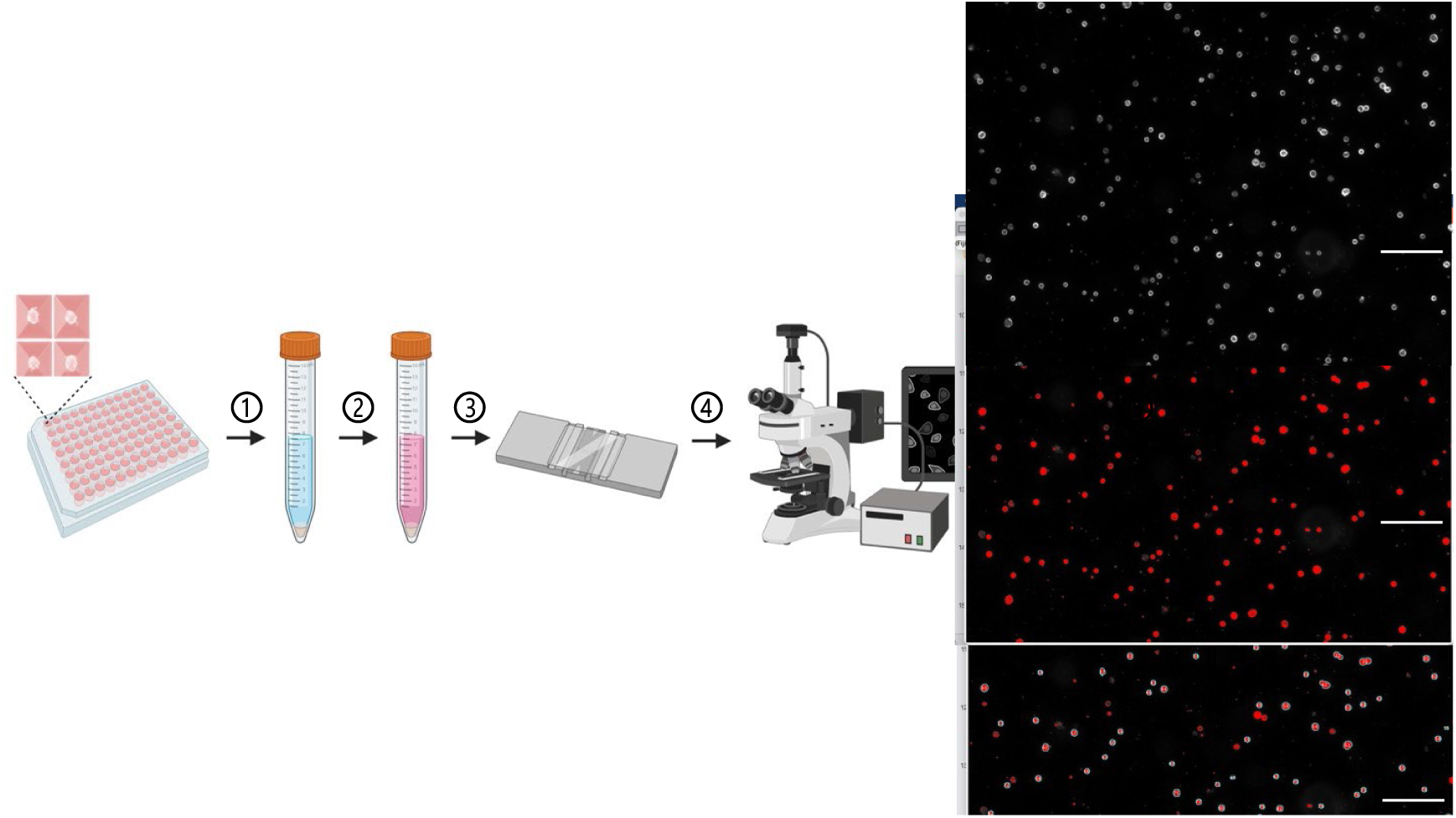
Cell size analysis procedure. ①: Harvest spheroids ②: Dissociate spheroids into single cells ③: Pipette the cell suspension into a hemocytometer ④: Image cells with a phase contrast microscopy ⑤: Analyze cell size using ImageJ a) Import phase contrast images to ImageJ b) Set scale using **Analyze – Set Scale** c) Adjust threshold using **Image – Adjust – Color Threshold** d) Measure cell area using **Analyze – Analyze Particles**, setting circularity to 0.70–1.00. e) Export results to Excel and calculate cell diameter using the formula: 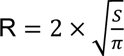 • R: Cell diameter • S: Area value from ImageJ

## References

1. Pittenger, M. F. et al. Mesenchymal stem cell perspective: cell biology to clinical progress. npj Regen. Med. 4, (2019).

2. Lim, M. et al. Intravenous injection of allogeneic umbilical cord-derived multipotent mesenchymal stromal cells reduces the infarct area and ameliorates cardiac function in a porcine model of acute myocardial infarction. Stem Cell Res. Ther. 9, 129 (2018).

3. Shake, J. G. et al. Mesenchymal stem cell implantation in a swine myocardial infarct model: Engraftment and functional effects. Ann. Thorac. Surg. 73, 1919–1926 (2002).

4. Berry, M. F. et al. Mesenchymal stem cell injection after myocardial infarction improves myocardial compliance. Am. J. Physiol. - Hear. Circ. Physiol. 290, H2196–H2203 (2006).

5. Luger, D. et al. Intravenously delivered mesenchymal stem cells: systemic anti-inflammatory effects improve left ventricular dysfunction in acute myocardial infarction and ischemic cardiomyopathy. Circ. Res. 120, 1598–1613 (2017).

6. Shafei, A. E. S. et al. Mesenchymal stem cell therapy: A promising cell-based therapy for treatment of myocardial infarction. J. Gene Med. 19, e2995 (2017).

7. Dong, F. et al. Myocardial CXCR4 expression is required for mesenchymal stem cell mediated repair following acute myocardial infarction. Circulation 126, 314–324 (2012).

8. Shyu, K. G., Wang, B. W., Hung, H. F., Chang, C. C. & Shih, D. T. B. Mesenchymal stem cells are superior to angiogenic growth factor genes for improving myocardial performance in the mouse model of acute myocardial infarction. J. Biomed. Sci. 13, 47–58 (2006).

9. Bagno, L., Hatzistergos, K. E., Balkan, W. & Hare, J. M. Mesenchymal stem cell-based therapy for cardiovascular disease: progress and challenges. Mol. Ther. 26, 1610–1623 (2018).

10. Yousefi, F., Ebtekar, M., Soleimani, M., Soudi, S. & Hashemi, S. M. Comparison of in vivo immunomodulatory effects of intravenous and intraperitoneal administration of adipose-tissue mesenchymal stem cells in experimental autoimmune encephalomyelitis (EAE). Int. Immunopharmacol. 17, 608–616 (2013).

11. Lei, F., Haque, R., Xiong, X. & Song, J. Modulation of autoimmune diseases by iPS cells. Methods Mol. Biol. 1213, 365–377 (2014).

12. Rafei, M., Birman, E., Forner, K. & Galipeau, J. Allogeneic mesenchymal stem cells for treatment of experimental autoimmune encephalomyelitis. Mol. Ther. 17, 1799–1803 (2009).

13. Wang, D., Li, S. P., Fu, J. S., Bai, L. & Guo, L. Resveratrol augments therapeutic efficiency of mouse bone marrow mesenchymal stem cell-based therapy in experimental autoimmune encephalomyelitis. Int. J. Dev. Neurosci. 49, 60–66 (2016).

14. Al Jumah, M. A. & Abumaree, M. H. The immunomodulatory and neuroprotective effects of mesenchymal stem cells (MSCs) in experimental autoimmune encephalomyelitis (EAE): A model of multiple sclerosis (MS). Int. J. Mol. Sci. 13, 9298–9331 (2012).

15. Morando, S. et al. The therapeutic effect of mesenchymal stem cell transplantation in experimental autoimmune encephalomyelitis is mediated by peripheral and central mechanisms. Stem Cell Res. Ther. 3, 3 (2012).

16. Chiang, E. R. et al. Allogeneic mesenchymal stem cells in combination with hyaluronic acid for the treatment of osteoarthritis in rabbits. PLoS One 11, e0149835 (2016).

17. Riester, S. M. et al. Safety studies for use of adipose tissue-derived mesenchymal stromal/stem cells in a rabbit model for osteoarthritis to support a phase I clinical trial. Stem Cells Transl. Med. 6, 910–922 (2017).

18. Deng, Z. et al. Human umbilical cord mesenchymal stem cells on treating osteoarthritis in a rabbit model: Injection strategies. Heliyon 10, e38384 (2024).

19. Kim, H. et al. Therapeutic effect of mesenchymal stem cells derived from human umbilical cord in rabbit temporomandibular joint model of osteoarthritis. Sci. Rep. 9, 13854 (2019).

20. Kim, S. E. et al. Intra-articular umbilical cord derived mesenchymal stem cell therapy for chronic elbow osteoarthritis in dogs: a double-blinded, placebo-controlled clinical trial. Front. Vet. Sci. 6, 474 (2019).

21. Maki, C. B. et al. Intra-articular Administration of Allogeneic Adipose Derived MSCs Reduces Pain and Lameness in Dogs With Hip Osteoarthritis: A Double Blinded, Randomized, Placebo Controlled Pilot Study. Front. Vet. Sci. 7, 1–11 (2020).

22. Zhang, B. Y. et al. Evaluation of the curative effect of umbilical cord mesenchymal stem cell therapy for knee arthritis in dogs using imaging technology. Stem Cells Int. 2018, 1983025 (2018).

23. Le Blanc, K. et al. Mesenchymal stem cells for treatment of steroid-resistant, severe, acute graft-versus-host disease: a phase II study. Lancet 371, 1579–1586 (2008).

24. Karussis, D. et al. Safety and immunological effects of mesenchymal stem cell transplantation in patients with multiple sclerosis and amyotrophic lateral sclerosis. Arch. Neurol. 67, 1187–1194 (2010).

25. He, X. et al. Umbilical cord-derived mesenchymal stem (stromal) cells for treatment of severe sepsis: aphase 1 clinical trial. Transl. Res. 199, 52–61 (2018).

26. McIntyre, L. A. et al. Cellular immunotherapy for septic shock: A phase I clinical trial. Am. J. Respir. Crit. Care Med. 197, 337–347 (2018).

27. Chen, J. et al. Clinical study of mesenchymal stem cell treatment for acute respiratory distress syndrome induced by epidemic influenza A (H7N9) infection: a hint for COVID-19 treatment. Engineering 6, 1153–1161 (2020).

28. Connick, P. et al. Autologous mesenchymal stem cells for the treatment of secondary progressive multiple sclerosis: An open-label phase 2a proof-of-concept study. Lancet Neurol. 11, 150–156 (2012).

29. Wilson, J. G. et al. Mesenchymal stem (stromal) cells for treatment of ARDS: A phase 1 clinical trial. Lancet Respir. Med. 3, 24–32 (2015).

30. Panés, J. et al. Expanded allogeneic adipose-derived mesenchymal stem cells (Cx601) for complex perianal fistulas in Crohn’s disease: a phase 3 randomised, double-blind controlled trial. Lancet. 388, 1281–1290 (2016).

31. Kebriaei, P. et al. A phase 3 randomized study of Remestemcel-L versus placebo added to second-line therapy in patients with steroid-refractory acute graft-versus-host disease. Biol. Blood Marrow Transplant. 26, 835–844 (2020).

32. Zheng, G. et al. Treatment of acute respiratory distress syndrome with allogeneic adipose-derived mesenchymal stem cells: a randomized, placebo-controlled pilot study. Respir. Res. 15, 39 (2014).

33. Lv, H. et al. Mesenchymal stromal cells as a salvage treatment for confirmed acute respiratory distress syndrome: preliminary data from a single-arm study. Intensive Care Med. 46, 1944–1947 (2020).

34. Matthay, M. A. et al. Treatment with allogeneic mesenchymal stromal cells for moderate to severe acute respiratory distress syndrome (START study): a randomised phase 2a safety trial. Lancet Respir. Med. 7, 154–162 (2019).

35. Gennadiy, G. et al. The results of the single center pilot randomized Russian clinical trial of mesenchymal stromal cells in severe neutropenic patients with septic shock (RUMCESS). Int. J. Blood Res. Disord. 5, 033 (2018).

36. Thompson, M. et al. Cell therapy with intravascular administration of mesenchymal stromal cells continues to appear safe: An updated systematic review and meta-analysis. EClinicalMedicine 19, 100249 (2020).

37. Mizukami, A. et al. Technologies for large-scale umbilical cord-derived MSC expansion: Experimental performance and cost of goods analysis. Biochem. Eng. J. 135, 36–48 (2018).

38. Mizukami, A. Mesenchymal Stromal Cells: From Discovery to Manufacturing and Commercialization. 2018, 4083921 (2018).

39. Roh, K. H., Nerem, R. M. & Roy, K. Biomanufacturing of Therapeutic Cells: State of the Art, Current Challenges, and Future Perspectives. Annu. Rev. Chem. Biomol. Eng. 7, 455–478 (2016).

40. Schnitzler, A. C. et al. Bioprocessing of human mesenchymal stem/stromal cells for therapeutic use: Current technologies and challenges. Biochem. Eng. J. 108, 3–13 (2016).

41. Skardal, A., Mack, D., Atala, A. & Sokern, S. Substrate elasticity controls cell proliferation, surface marker expression and motile phenotype in amniotic fluid-derived stem cells. J. Mech. Behav. Biomed. Mater. 17, 307–316 (2013).

42. Huebsch, N. Translational mechanobiology: Designing synthetic hydrogel matrices for improved in vitro models and cell-based therapies. Acta Biomater. 94, 97–111 (2019).

43. Yang, Y. H. K., Ogando, C. R., Wang See, C., Chang, T. Y. & Barabino, G. A. Changes in phenotype and differentiation potential of human mesenchymal stem cells aging in vitro. Stem Cell Res. Ther. 9, 131 (2018).

44. Kureel, S. K. et al. Soft substrate maintains proliferative and adipogenic differentiation potential of human mesenchymal stem cells on long-term expansion by delaying senescence. Biol. Open 8, bio039453 (2019).

45. Chen, Y. et al. Mesenchymal Stem Cell: Considerations for Manufacturing and Clinical Trials on Cell Therapy Product ClinMed. *Int*. J. Stem Cell Res. Ther. 3, 029 (2016).

46. Weiss, D. J. Cell-based therapies for acute respiratory distress syndrome. Lancet Respir. Med. 7, 105–106 (2019).

47. Galipeau, J. The mesenchymal stromal cells dilemma-does a negative phase III trial of random donor mesenchymal stromal cells in steroid-resistant graft-versus-host disease represent a death knell or a bump in the road? Cytotherapy 15, 2–8 (2013).

48. Mo, M., Zhou, Y., Li, S. & Wu, Y. Three-Dimensional Culture Reduces Cell Size By Increasing Vesicle Excretion. Stem Cells 36, 286–292 (2018).

49. Ge, J. et al. The size of mesenchymal stem cells is a significant cause of vascular obstructions and stroke. Stem Cell Rev. Rep. 10, 295–303 (2014).

50. Lu, R. et al. 3D spheroid culture synchronizes heterogeneous MSCs into an immunomodulatory phenotype with enhanced anti-inflammatory effects. iScience 27, 110811 (2024).

51. Ji, X. et al. Impact of mesenchymal stem cell size and adhesion modulation on in vivo distribution: insights from quantitative PET imaging. Stem Cell Res. Ther. 15, 456 (2024).

52. Zhuang, W. Z. et al. Mesenchymal stem/stromal cell-based therapy: mechanism, systemic safety and biodistribution for precision clinical applications. J. Biomed. Sci. 28, 28 (2021).

53. Sanchez-Diaz, M. et al. Biodistribution of mesenchymal stromal cells after administration in animal models and humans: A systematic review. J. Clin. Med. 10, 2925 (2021).

54. Guo, L. et al. Three-dimensional spheroid-cultured mesenchymal stem cells devoid of embolism attenuate brain stroke injury after intra-arterial injection. Stem Cells Dev. 23, 978–989 (2014).

55. Janowski, M. et al. Cell size and velocity of injection are major determinants of the safety of intracarotid stem cell transplantation. J. Cereb. Blood Flow Metab. 33, 921–927 (2013).

56. Cesarz, Z. & Tamama, K. Spheroid Culture of Mesenchymal Stem Cells. Stem Cells Int. 2016, 9176357 (2016).

57. Hazrati, A., Malekpour, K., Soudi, S. & Hashemi, S. M. Mesenchymal stromal/stem cells spheroid culture effect on the therapeutic efficacy of these cells and their exosomes: A new strategy to overcome cell therapy limitations. Biomed. Pharmacother. 152, 113211 (2022).

58. Ohori-Morita, Y., Ashry, A., Niibe, K. & Egusa, H. Current perspectives on the dynamic culture of mesenchymal stromal/stem cell spheroids. Stem Cells Transl. Med. 14, szae093 (2025).

59. Bijonowski, B. M. et al. Aggregation-induced integrated stress response rejuvenates culture-expanded human mesenchymal stem cells. Biotechnol. Bioeng. 117, 3136–3149 (2020).

60. Bijonowski, B. M., Yuan, X., Jeske, R., Li, Y. & Grant, S. C. Cyclical aggregation extends in vitro expansion potential of human mesenchymal stem cells. Sci. Rep. 10, 20448 (2020).

61. Han, L., et al. Mesenchymal stromal cells and alpha-1 antitrypsin have a strong synergy in modulating inflammation and its resolution. Theranostics 13, 2843–2862 (2023).

62. Wang, O. & Lei, Y. Creating a cell-friendly microenvironment to enhance cell culture efficiency. Cell Gene Ther. Insights 5, 341–350 (2019).

63. Wang, Z. et al. Comparative Study of Human Pluripotent Stem Cell-Derived Endothelial Cells in Hydrogel-Based Culture Systems. ACS Omega 6, 6942–6952 (2021).

64. Lin, H. et al. Manufacturing human pluripotent stem cell derived endothelial cells in scalable and cell-friendly microenvironments. Biomater. Sci. 7, 373–388 (2019).

65. Lin, H. et al. Integrated generation of induced pluripotent stem cells in a low-cost device. Biomaterials 189, 23–36 (2019).

66. Yang, Y., et al. Collagen Hydrogel Tube Microbioreactors for Cell and Tissue Manufacturing. bioRxiv (2025).

67. Lin, H. et al. Manufacturing human pluripotent stem cell derived endothelial cells in scalable and cell-friendly microenvironments. Biomater. Sci. 7, 373–388 (2019).

68. Li, Q. et al. Scalable culturing of primary human glioblastoma tumor-initiating cells with a cell-friendly culture system. Sci. Rep. 8, 3531 (2018).

69. Lin, H. et al. Hydrogel-Based Bioprocess for Scalable Manufacturing of Human Pluripotent Stem Cell-Derived Neural Stem Cells. ACS Appl. Mater. Interfaces 10, 29238–29250 (2018).

70. Lin, H. et al. Engineered Microenvironment for Manufacturing Human Pluripotent Stem Cell-Derived Vascular Smooth Muscle Cells. Stem Cell Reports 12, 84–97 (2019).

71. Lin, H. et al. Engineered Microenvironment for Manufacturing Human Pluripotent Stem Cell-Derived Vascular Smooth Muscle Cells. Stem Cell Reports 12, 84–97 (2019).

72. Liu, Q. et al. Comparative study of differentiating human pluripotent stem cells into vascular smooth muscle cells in hydrogel-based culture methods. Regen. Ther. 22, 39–49 (2023).

73. Li, Q. et al. Scalable and physiologically relevant microenvironments for human pluripotent stem cell expansion and differentiation. Biofabrication 10, 025006 (2018).

74. Lin, H. et al. Automated expansion of primary human T cells in scalable and cell-friendly hydrogel microtubes for adoptive immunotherapy. Adv. Healthc. Mater. 7, e1701297 (2018).

75. Binato, R. et al. Stability of human mesenchymal stem cells during in vitro culture: Considerations for cell therapy. Cell Prolif. 46, 10–22 (2013).

76. Gu, Y. et al. Changes in mesenchymal stem cells following long-term culture in vitro. Mol. Med. Rep. 13, 5207–5215 (2016).

77. Weng, Z. et al. Mesenchymal Stem/Stromal Cell Senescence: Hallmarks, Mechanisms, and Combating Strategies. Stem Cells Transl. Med. 11, 356–371 (2022).

78. Roger, L., Tomas, F. & Gire, V. Mechanisms and regulation of cellular senescence. Int. J. Mol. Sci. 22, 13173 (2021).

79. Neri, S. & Borzì, R. M. Molecular mechanisms contributing to mesenchymal stromal cell aging. Biomolecules 10, 340 (2020).

80. Zhou, X., Hong, Y., Zhang, H. & Li, X. Mesenchymal Stem Cell Senescence and Rejuvenation: Current Status and Challenges. Front. Cell Dev. Biol. 8, 364 (2020).

81. Kouroupis, D. & Correa, D. Increased Mesenchymal Stem Cell Functionalization in Three-Dimensional Manufacturing Settings for Enhanced Therapeutic Applications. Front. Bioeng. Biotechnol. 9, 621748 (2021).

82. Duval, K. et al. Modeling physiological events in 2D vs. 3D cell culture. Physiology 32, 266–277 (2017).

83. Lengefeld, J. et al. Cell size is a determinant of stem cell potential during aging. Sci. Adv. 7, eabk0271 (2021).

84. Neurohr, G. E. et al. Excessive Cell Growth Causes Cytoplasm Dilution And Contributes to Senescence. Cell 176, 1083–1097.e18 (2019).

85. Sun, Y. et al. Spheroid-cultured human umbilical cord-derived mesenchymal stem cells attenuate hepatic ischemia-reperfusion injury in rats. Sci. Rep. 8, 2518 (2018).

86. Fuentes, P. et al. Dynamic Culture of Mesenchymal Stromal/Stem Cell Spheroids and Secretion of Paracrine Factors. Front. Bioeng. Biotechnol. 10, 916229 (2022).

87. Cesarz, Z. & Tamama, K. Spheroid Culture of Mesenchymal Stem Cells. Stem Cells Int. 2016, 9176357(2016).

88. Vanover, M. et al. High density placental mesenchymal stromal cells provide neuronal preservation and improve motor function following in utero treatment of ovine myelomeningocele. J. Pediatr. Surg. 54, 75–79 (2019).

89. Yamashiro, K. et al. Surviving Lambs with Myelomeningocele Repaired in utero with Placental Mesenchymal Stromal Cells for 6 Months: A Pilot Study. Fetal Diagn. Ther. 47, 912–917 (2020).

90. Stokes, S. C. et al. Impact of Gestational Age on Neuroprotective Function of Placenta-Derived Mesenchymal Stromal Cells. J. Surg. Res. 273, 201–210 (2022).

91. Jackson, J. E. et al. Placental Mesenchymal Stromal Cells: Preclinical Safety Evaluation for Fetal Myelomeningocele Repair. J. Surg. Res. 267, 660–668 (2021).

92. Kabagambe, S. et al. Placental mesenchymal stromal cells seeded on clinical grade extracellular matrix improve ambulation in ovine myelomeningocele. J. Pediatr. Surg. 53, 178–182 (2018).

93. Theodorou, C. M. et al. Early investigations into improving bowel and bladder function in fetal ovine myelomeningocele repair. J. Pediatr. Surg. 57, 941–948 (2022).

94. Clark, K. C. et al. Translational applications of placental dervided mesenchymal stem cells for the treatment of spina bifida: a canine model. Cytotherapy 21, S75 (2019).

95. Stokes, S. C. et al. 154 Placental MSCs for in utero myelomeningocele repair do not present additional risk for pregnant ewes. Am. J. Obstet. Gynecol. 224, S106 (2021).

96. Pan, W., Chen, H., Wang, A., Wang, F. & Zhang, X. Challenges and strategies: Scalable and efficient production of mesenchymal stem cells-derived exosomes for cell-free therapy. Life Sci. 319, 121524 (2023).

97. Chen, Y. J. et al. Fetal surgical repair with placenta-derived mesenchymal stromal cell engineered patch in a rodent model of myelomeningocele. J. Pediatr. Surg. 53, 183–188 (2018).

98. Theodorou, C. M. et al. Efficacy of clinical-grade human placental mesenchymal stromal cells in fetal ovine myelomeningocele repair. J. Pediatr. Surg. 57, 753–758 (2022).

99. Zhang, X. et al. Engineered neuron-targeting, placental mesenchymal stromal cell-derived extracellular vesicles for in utero treatment of myelomeningocele. bioRxiv (2021).

100. Stokes, S. C. et al. Long-term safety evaluation of placental mesenchymal stromal cells for in utero repair of myelomeningocele in a novel ovine model. J. Pediatr. Surg. 57, 18–25 (2022).

101. Long, C., Lankford, L. & Wang, A. Stem cell-based in utero therapies for spina bifida: Implications for neural regeneration. Neural Regen. Res. 14, 260–261 (2019).

102. Prins Henk-Jan, S. E. B. C. Placental Mesenchymal Stromal Cells Rescue Ambulation in Ovine Myelomeningocele. Stem Cells Transl. Med. 4, 659–669 (2014).

103. Lankford, L. et al. Manufacture and preparation of human placenta-derived mesenchymal stromal cells for local tissue delivery. Cytotherapy 19, 680–688 (2017).

104. Galganski, L. A. et al. In utero treatment of myelomeningocele with placental mesenchymal stromal cells — Selection of an optimal cell line in preparation for clinical trials. J. Pediatr. Surg. 55, 1941–1946 (2020).

